# Reducing type II error in fMRI analysis: Cluster-extent threshold simulation results and an evaluation of current methods to correct for multiple comparisons

**DOI:** 10.1101/2025.07.23.666419

**Authors:** Scott D. Slotnick

## Abstract

Many procedures to correct for multiple comparisons in functional magnetic resonance imaging (fMRI) analysis require a minimum cluster-extent threshold; however, sample size (N) is often not modeled. In this study, a series of simulations was conducted where N was varied to determine whether this parameter affected cluster threshold. The primary hypothesis was that modeling N in the simulations would reduce cluster thresholds. A secondary hypothesis was that this cluster size reduction was due to between-subject variability, which was tested by eliminating the corresponding standard error term. Acquisition volume parameters were fixed, while key parameters were varied to reflect reasonable ranges: N (10, 20, or 30), corrected p-value (.05, .01, or .001), individual-voxel p-value (.01, .005, or .001), FWHM (3, 5, or 7 mm), and voxel resolution (2 or 3 mm). Each simulation consisted of 100 iterations repeated 100 times, with a total of 4,860,000 iterations and 66,420,000 simulated subjects. There was a significant effect of condition with clusters approximately 18% smaller with versus without N modeled and a significant increase in cluster thresholds for larger sample sizes. Bayesian analysis provided very strong support for the secondary hypothesis. These simulation results were replicated in a real fMRI data set. The present findings indicate that sample size should be incorporated into all methods to provide the most accurate thresholds possible and reduce type II error. A broader range of topics is discussed including balancing type I and type II error, and the assumption that non-task fMRI activity reflects null data is questioned.

---

Analysis of functional magnetic resonance imaging (fMRI) data typically involves conducting statistical tests for hundreds of thousands of voxels in the brain volume such that it is necessary to correct for multiple comparisons. There are multiple methods to correct for multiple comparisons in fMRI analysis. Bonferroni correction requires dividing the desired corrected p-value (i.e., the familywise type I error rate) by the total number of comparisons, but this is method is too strict for a whole-brain analysis because of the high rate of type II error (i.e., failure to detect real effects). With the aim of balancing type I and type II error, Benjamini and Hochberg (1995) derived a procedure to limit the false discovery rate (i.e., the ratio of false activations to the sum of true and false activations; see also, Benjamini et al., 2006). In the original version of the analysis software AFNI (Cox, 1996), Monte Carlo simulations were conducted with brain activity in the acquisition volume modeled by random values with Gaussian spatial autocorrelation, and the minimum spatial extent of contiguous activity across simulations was computed to correct for multiple comparisons (as larger clusters of spurious activity are less likely by chance). Slotnick et al. (2003) developed a stand-alone script in MATLAB (Natick, MA), cluster_threshold_beta (Slotnick, 2025a), that used a similar cluster-extent correction procedure (see the Methods). A limitation of these cluster-extent threshold procedures is that they do not implement sample size. The present findings will show that this may result in increased type II error due to overly strict (large) cluster thresholds.

Other methods to correct for multiple comparisons use actual brain activity, rather than simulated brain activity, to correct for multiple comparisons. The analysis software SPM assumes the group residual activation map, which remains after all events of interest are modeled, along with Gaussian random field theory (Kiebel et al., 1999), to conduct cluster-extent correction for multiple comparisons. The analysis software FSL employes the same method (Jenkinson et al., 2012). Eklund et al. (2016) assumed that resting-state (non-task) data reflected null data (see the Discussion for an evaluation of this assumption), and they compared familywise type I error rates for AFNI, SPM, FSL, and their own (non-cluster-extent) nonparametric method, and presented evidence indicating that all cluster-extent correction methods had unacceptable false-positive rates. In a Discussion paper (Slotnick, 2017a), I argued Eklund et al.’s assumption that resting-state data reflected null data was incorrect because the corresponding activation maps showed robust activation in default-network regions that falsely inflated type I error, while true null data should reflect noise and be evenly distributed across the brain (for further discussion, see Nichols et al, 2017; Slotnick, 2017b). Seemingly motivated by Eklund et al. (2016), AFNI implemented a new cluster-extent correction method that assumes the spatial-autocorrelation function has a ‘long tail’ (deviating from the previous Gaussian assumption; Cox, 2017), which is the currently recommended method to correct for multiple comparisons for that software (AFNI program, 2025a). Of importance, the use of the residual brain activation map in SPM and FSL, which reflects non-task activity, may similarly include true activations from default-network regions that will yield larger cluster-extent thresholds than if true null data volumes were employed. As such, the methods commonly employed in these analysis software programs (AFNI, SPM, and FSL) could be overly conservative and yield inflated type II error rates (see the Discussion for a more thorough assessment of this issue).

Although the large majority of fMRI users seem primarily concerned with type I error, compelling arguments have been made that we should balance type I and type II error (Liberman & Cunningham, 2009; Hopfinger, 2017). That is, focusing on only type I error can result in a number of negative consequences including increasing type II error, a bias toward publishing obvious results, deficient meta-analyses, and stifling innovation (see the Discussion for consideration of these and related points in more detail). Rather than using stricter thresholds and focusing on only type I error, we should use optimal methods to balance type I and type II error.

In the present study, with the aim of potentially reducing type II error, in a series of Monte Carlo simulation sets, sample size was parametrically varied to determine whether this parameter affected cluster threshold using a new script called cluster_threshold_gamma (Slotnick, 2025b). The primary hypothesis was that incorporating sample size into the simulations would reduce the cluster-extent threshold, as the magnitude of activity in voxels at cluster boundaries was expected to be particularly variable across subjects (such that activity in these voxels would drop out, thereby reducing cluster size). A secondary hypothesis was that cluster thresholds estimated by cluster_threshold_gamma would systematically increase for larger sample sizes, which are associated with lower standard errors and thus higher t-values.

## Simulation Set 1

### Methods

#### fMRI simulations

Simulations were conducted using the cluster_threshold_beta script (Slotnick, 2025a) and the cluster_threshold_gamma script (Slotnick, 2025b), which were written in MATLAB and are published on the Open Science Framework (https://osf.io/). It is notable that the first (unreleased) version of the cluster_threshold script was called cluster_threshold_alpha, thus cluster_threshold_beta and cluster_threshold_gamma were named in sequence following the Greek alphabet (and cluster_threshold_beta is actually a triple entendre, also alluding to a software beta release and fMRI beta weights). In brief, cluster_threshold_beta is a function where a user inputs fMRI acquisition volume dimensions (x_matrix, y_matrix, slices), voxel dimensions (in mm; dim_xy, dim_z), the size of the full-width-half-maximum (FWHM) smoothing kernel (in mm), resampled voxel dimensions (in mm), desired corrected p-value, specified individual-voxel p-value, and the number of Monte Carlo simulations. The critical output is the minimum number of contiguous resampled voxels that are required to correct for multiple comparisons, which is referred to as the cluster-extent threshold or cluster threshold. Within the function, brain activity is modeled at each voxel (i.e., independently) within the acquisition volume using a normally distributed random number with a mean of 0 and a standard deviation of 1 (with 99.7% of the data having values between –3 and 3). Based on the central limit theorem, it is reasonable to assume that voxel activation magnitudes have a normal distribution. Across the entire volume, this activity is smoothed using a three-dimensional gaussian kernel (i.e., the specified FWHM size) to model spatial autocorrelation, the activity is thresholded to produce significant activations corresponding to the individual-voxel p-value employed (e.g., at a p-value of .001, across the entire brain volume, the threshold is adjusted such that approximately 1 out of every 1,000 voxels will significant), the size of all contiguous clusters of activity are counted in the brain volume, and this process is repeated according to the number of specified iterations (typically 10,000). The cluster sizes are summed across all iterations, and the cluster threshold is determined where that cluster size or larger occurs less than the desired corrected p-value. Cluster_threshold_gamma is identical except that the number of subjects (N) is specified as an input parameter, brain activity for each subject is modeled using normally distributed random numbers, and then this activity is smoothed for each subject using the specified gaussian kernel. A group t-map null volume is created where the t-value at each voxel is computed by dividing the between-subject mean magnitude of activity by the standard error (i.e., a random-effect analysis is conducted). This t-map is thresholded to produce the specified individual-voxel p-value and the remaining steps are identical to those described above for cluster_threshold_beta. Note that cluster_threshold_beta and cluster_threshold_gamma were intended to provide the cluster-extent threshold for a standard univariate general linear model analysis.

For the simulations conducted, acquisition volume parameters were fixed with x_matrix = 64, y_matrix = 64, slices = 45, dim_xy = 3 mm, dim_z = 3 mm, and dim_resampled = 2 mm to simulate a standard whole-brain acquisition and analysis protocol. Other key input parameters were parametrically varied to reflect reasonable ranges including sample size (N = 10, 20, 30), corrected p-value (.05, .01, .001), individual-voxel p-value (.01, .005, .001), and FWHM (3, 5, 7). The FWHM values were selected with the aim of simulating null activation volumes with the following considerations: 1) in my experience, the minimum spatial autocorrelation in fMRI group activation maps with very few activations is approximately 3 mm (e.g., Spets & Slotnick, 2019; Spets et al., 2021a, 2021b), which creates a lower limit for this parameter, 2) null volumes should have relatively small spatial autocorrelation values thus motivating increasing this parameter by relatively small increments of 2 mm, and 3) these values, used to model null data, were exactly half those used in a classic study (Hopfinger et al., 2000) that evaluated the effect of analysis parameters on a real fMRI data set, which is expected to have larger spatial autocorrelation values than null data (i.e., Hopfinger et al. employed FWHM values of 6, 10, and 14 mm). Although N was not a parameter for cluster_threshold_beta, new simulations were conducted for each of the values such that all data points were independent. For a given input parameter set, each Monte Carlo simulation consisted of 100 iterations resulting in a single cluster-extent threshold and each simulation was repeated 100 times to provide a measure of variance and allow for statistical analysis. Of importance, this translates to each data point in the figures reflecting 10,000 iterations. It is notable that for the two conditions (cluster_threshold_beta, cluster_threshold_gamma), there were a total of 1,620,000 iterations with 17,010,000 simulated subjects (assuming each cluster_threshold_beta iteration consisted of a single subject). These stimulations were conducted using the Boston College Linux High-Performance Computing (HPC) cluster, simultaneously running 400 jobs, each with 1 CPU and 8 GB of memory, for approximately 1 week.

To illustrate the primary results, for each condition (cluster_threshold_beta, cluster_threshold_gamma) and each parameter set, the mean cluster threshold (for 100 iterations) was plotted with 95% confidence intervals. To quantify cluster threshold sizes between conditions, the ratio of cluster threshold means was also computed along with 95% confidence intervals, assuming standard error propagation theory of independent normally distributed random variables (Holmes & Buhr, 2007).

### Statistical analysis

Statistical analysis was conducted using JASP (JASP Team, 2024). A frequentist two-factor ANOVA was used to test the primary hypothesis that the cluster threshold (the dependent variable) was modulated by condition (cluster_threshold_beta, cluster_threshold_gamma) and also assessed whether there was a condition x sample size interaction. A Bayesian ANOVA with the same factors was used to determine the strength of evidence in support of the null hypothesis or the alternative hypothesis (Wagenmakers et al., 2018). Bayes factor values were interpreted following Kass and Raftery (1995), where –2log(BF_10_) values from 2–6 indicate positive evidence, values from 6–10 indicate strong evidence, and values greater than 10 indicate very strong evidence. For completeness, a series of two-factor ANOVAs was also conducted to confirm that cluster threshold was modulated by corrected p-value, individual-voxel p-value, and FWHM, along with exploratory analyses to determine whether condition interacted with each of these factors. Given that 40 independent statistical tests were conducted across three simulation sets, a single test was deemed significant if the p-value was less than .00125 (i.e., .05/40; Bonferroni corrected for multiple comparison).

### Results

Figure 1 shows the cluster-extent thresholds for each condition with FWHM smoothing kernel values of 3 mm, 5 mm or 7 mm. It is notable the 95% confidence intervals were very small (often zero in magnitude), illustrating that the cluster-extent thresholds were very stable, even with only 100 iterations. Of direct relevance to the primary hypothesis under investigation, there was a clear difference between conditions, with cluster_threshold_gamma (in red) producing smaller thresholds than cluster_threshold_beta (in blue) for most of the comparisons. As expected, cluster_threshold_beta thresholds did not change as a function of N; however, cluster_threshold_gamma thresholds usually increased as a function of N. A frequentist analysis confirmed that there was a significant main effect of condition (cluster_threshold_beta vs. cluster_threshold_gamma, F(1,16194) = 205.180, p < .001, η^2^ = .012), a significant main effect of N (F(2,16194) = 20.903, p < .001, η^2^ = .003), and a significant condition x N interaction (F(2,16194) = 20.601, p < .001, η^2^ = .002). A Bayesian analysis supported these findings as the interaction model (with Condition + N + Condition x N) had a posterior model probability close to unity (P(M) = 1.000), and all other models, including the null model, had Bayes factor values close to zero (all BF_10_ values < 8.605x10^-7^), which is very strong evidence against the null model. Follow-up analyses showed that cluster size was modulated by N in the cluster_threshold_gamma condition (frequentist, F(2,8097) = 50.313, p < .001, η^2^ = .012; Bayesian, N model P(M) = 1.00, null model BF_10_ = 1.584x10^-19^, which is very strong evidence against the null model), but was not modulated by N in the cluster_threshold_beta condition (frequentist, F(2,8097) = .005, p = .995, η^2^ = 1.299x10^-6^; Bayesian, null model P(M) = .999, N model BF_10_ = 0.001, which is very strong evidence favoring the null model).

**Figure 1.**
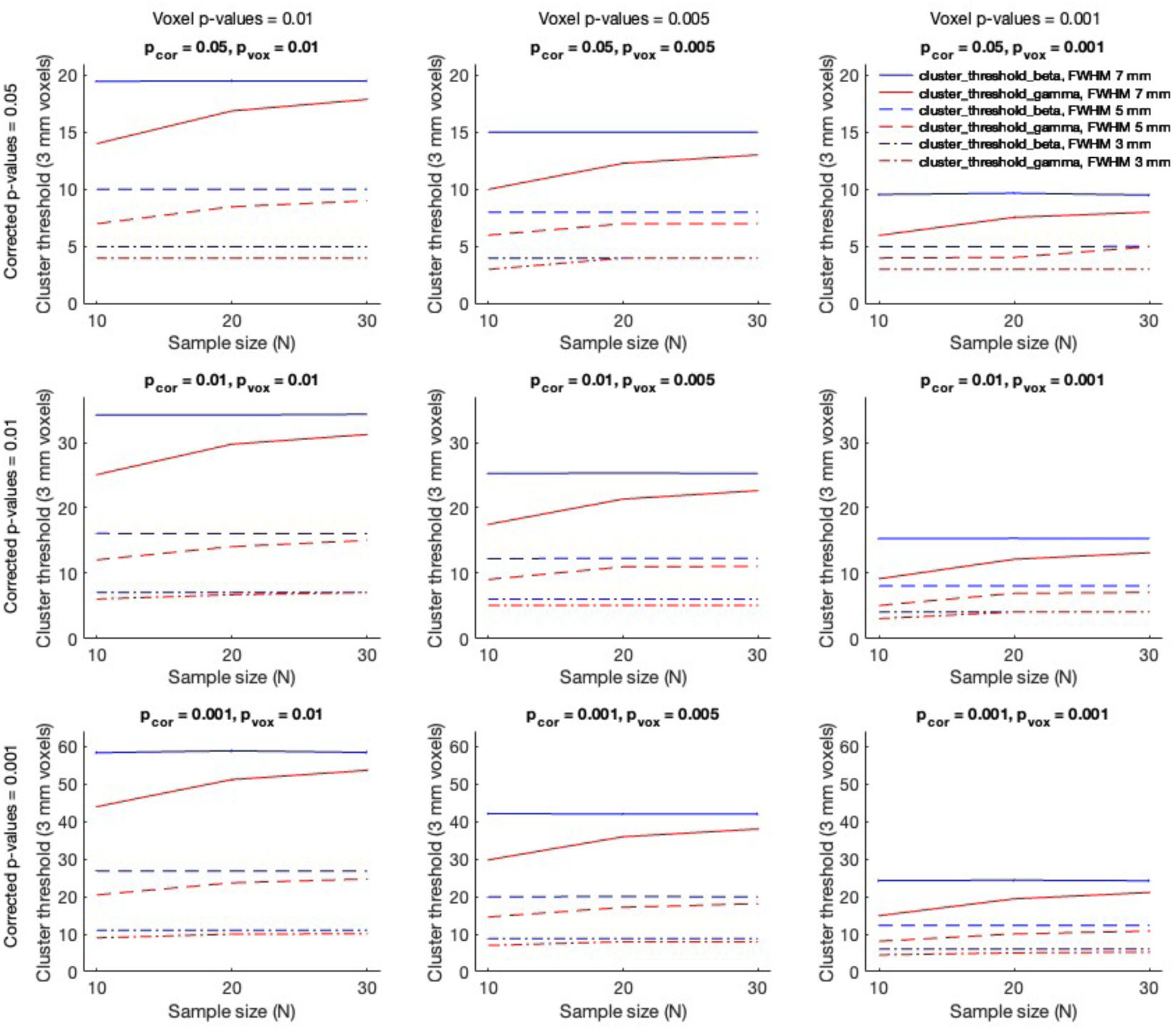
Cluster-extent thresholds (3 mm isotropic voxels) produced by cluster_threshold_beta (in blue) and cluster_threshold_gamma (in red) as a function of corrected p-value (p_cor) and individual-voxel p-value (p_vox) with a full-width-half-maximum (FWHM) smoothing kernel of 3 mm, 5 mm, or 7 mm (key at the top right). In each panel, for each simulation set (N = 10, 20, and 30), the mean and 95% confidence intervals are shown, with lines connecting the means for each condition to illustrate trends as a function of N.

As illustrated in Figure 1, as expected, there also appeared to be greater differential condition effects for stricter corrected p-values (which corresponds to larger cluster thresholds) and greater differential condition effects for stricter individual-voxel p-values (which corresponds to smaller cluster thresholds). For corrected p-values, there was a significant main effect of corrected p-value (F(2,16194) = 2233.825, p < .001, η^2^ = .213), and a significant condition x corrected p-value interaction (F(2,16194) = 18.289, p < .001, η^2^ = .002; this is illustrated by parametrically increasing condition effects from top to bottom in the panels of Figure 1). Bayesian analysis supported these findings as the interaction model (with condition + corrected p-value + condition x corrected p-value) had a posterior model probability close to unity (P(M) = 1.000) and all other models had Bayes factor values close to zero (all BF_10_ values < 8.533x10^-6^), which is very strong evidence against the null model. For individual-voxel p-values, there was a significant main effect of individual-voxel p-value (F(2,16194) = 1261.115, p < .001, η^2^ = .133), but the condition x individual-voxel p-value interaction was not significant (F(2,16194) = 3.537, p < .029, η^2^ = 3.731x10^-4^; this is illustrated by the relatively similar condition effects from left to right in the panels of Figures 1). Bayesian analysis supported these findings as the condition + individual-voxel p-value model had a very high posterior model probability (P(M) = .954), the interaction model (with condition + individual-voxel p-value + condition x individual-voxel p-value) had the second highest posterior model probability (P(M) = .046), the sum of these posterior probabilities was close to unity (P(M) = 1.000), and all other models had Bayes factor values close to zero (all BF_10_ values < 8.576x10^-50^), which is very strong evidence against the null model.

Figures 1 also appeared to show the expected greater differential condition effects with increasing FWHM smoothing kernels (which correspond to larger cluster thresholds). As a function of FWHM, there was a significant main effect of FWHM (F(2,16194) = 6384.941, p < .001, η^2^ = .433), and a significant condition x FWHM interaction (F(2,16194) = 83.042, p < .001, η^2^ = .006). Bayesian analysis supported these findings as the interaction model (with condition + FWHM + condition x FWHM) had a posterior model probability close to unity (P(M) = 1.000) and all other models had Bayes factor values close to zero (all BF_10_ values < 1.207x10^-33^), which is very strong evidence against the null model.

As mentioned above, cluster thresholds produced by cluster_threshold_gamma were usually smaller than those produced by cluster_threshold_beta. To quantify this effect, the cluster threshold produced by cluster_threshold_gamma was divided by the cluster threshold produced by cluster_threshold_beta to produce a ratio. These results are illustrated in Figure 2 for 3, 5, and 7 mm FWHM smoothing kernels. This ratio ranged from 0.5968 to 1.0000, had a mean of 0.8391, and had a median of 0.8570. Thus, the average cluster threshold size produced by cluster_threshold_gamma was 16.09% smaller than the average cluster threshold size produced by cluster_threshold_beta.

**Figure 2.**
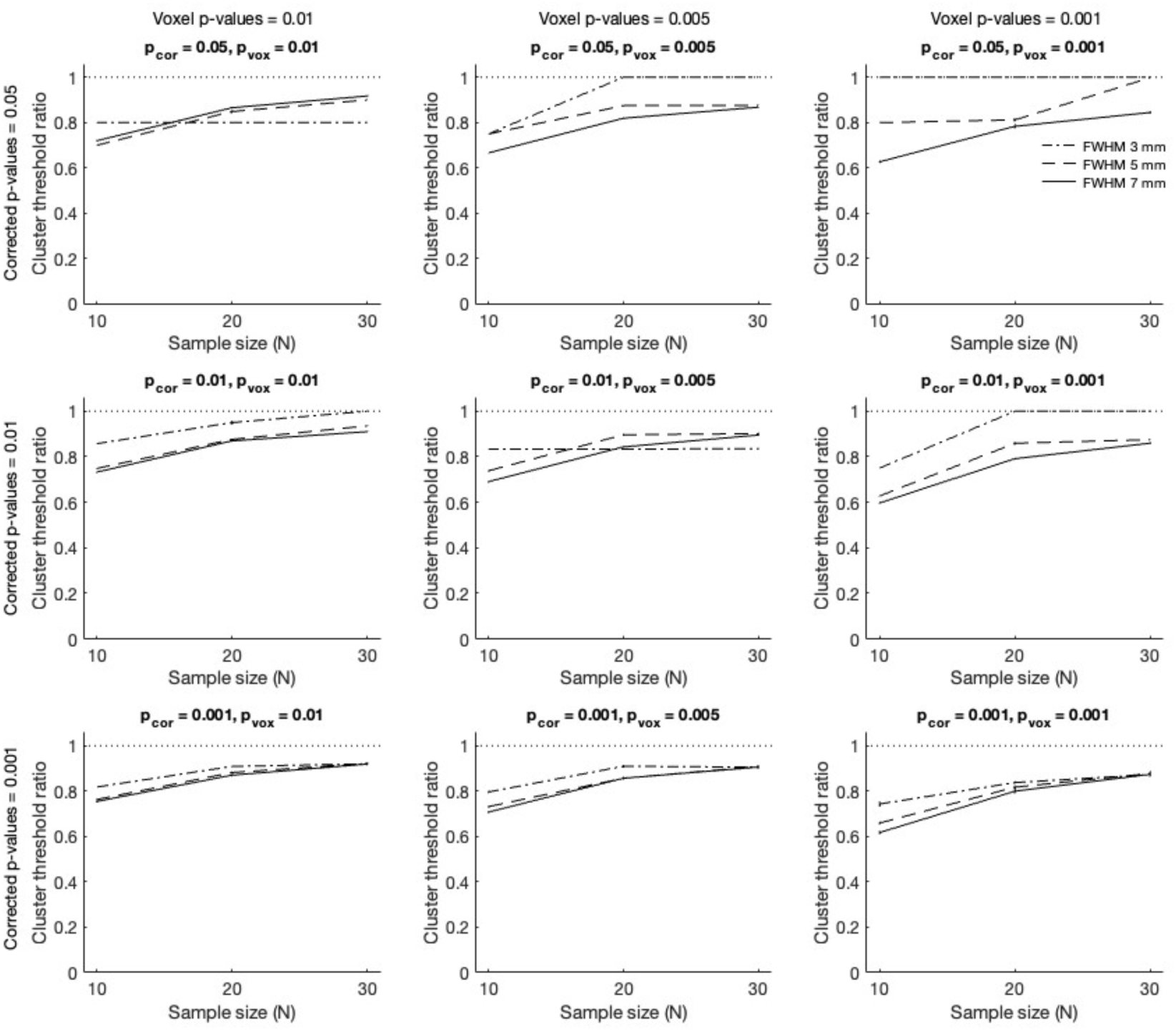
Ratio of cluster thresholds (3 mm isotropic voxels) produced by cluster_threshold_gamma and cluster_threshold_beta as a function of corrected p-value (p_cor) and individual-voxel p-value (p_vox) with a FWHM smoothing kernel of 3 mm, 5 mm, or 7 mm (key at the top right; the dotted line demarcates a ratio of 1, for reference). In each panel, for each simulation set (N = 10, 20, and 30), the ratio of means and 95% confidence intervals are shown, with lines connecting the means for each condition to illustrate trends as a function of N.

## Simulation Set 2

The previous results provided evidence that cluster thresholds produced with cluster_threshold_gamma were often smaller than those produced by cluster_threshold_beta. For the cluster_threshold_gamma simulations, this may have been due to activations near the outer border of clusters being less consistent across subjects (reflected by larger standard errors in the corresponding voxels) that would decrease the likelihood that these activations would be significant in the random-effect group analysis and thus reduce cluster threshold sizes. To test this hypothesis, a new set of simulations was conducted by slightly altering the cluster_threshold_gamma script, with the only change being that the between-subject mean value was not divided by the standard error (i.e., voxel activity was simulated by the between-subject mean value rather than the corresponding t-value). These simulations are subsequently referred to as cluster_threshold_nose (cluster_threshold_gamma with no between-subject standard error). If the hypothesis is correct, cluster thresholds produced by cluster_threshold_nose should not significantly differ from those produced by cluster_threshold_beta, which was tested in this simulation set.

### Methods

Unless otherwise specified, this simulations used the identical procedures as Simulation Set 1. For the new cluster_threshold_nose condition, there were 810,000 iterations with 16,200,000 simulated subjects. These stimulations were conducted using the Boston College Linux HPC, simultaneously running 350 jobs for approximately 1 week. Cluster_threshold_beta values were taken from the previous simulation set.

### Results

Figure 3 shows the cluster-extent thresholds for each condition with FWHM smoothing kernel values of 3 mm, 5 mm, or 7 mm. In support of the preceding hypothesis, there was no clear difference between conditions, with cluster_threshold_nose (in green) producing identical or nearly identical thresholds as cluster_threshold_beta (in blue) for nearly all comparisons (note, green lines with no hint of blue indicate perfect overlap). Thresholds did not appear to change as a function of N for cluster_threshold_nose, which is expected given that between-subject standard error values no longer modulated cluster size. A frequentist analysis confirmed that there was no significant main effect of condition (cluster_threshold_beta vs. cluster_threshold_nose, F(1,16194) = 9.428x10^-6^, p =.998, η^2^ = 5.822x10^-10^), no significant main effect of N (F(2,16194) = 6.498x10^-4^, p = .999, η^2^ = 8.025x10^-8^), and no significant condition x N interaction (F(2,16194) = .006, p = .994, η^2^ = 7.336x10^-7^). A Bayesian analysis supported these findings as the null model had a posterior model probability close to unity (P(M) = .982) and all other models had Bayes factor values close to zero (all BF_10_ values < 0.019), which is strong evidence favoring the null model.

**Figure 3.**
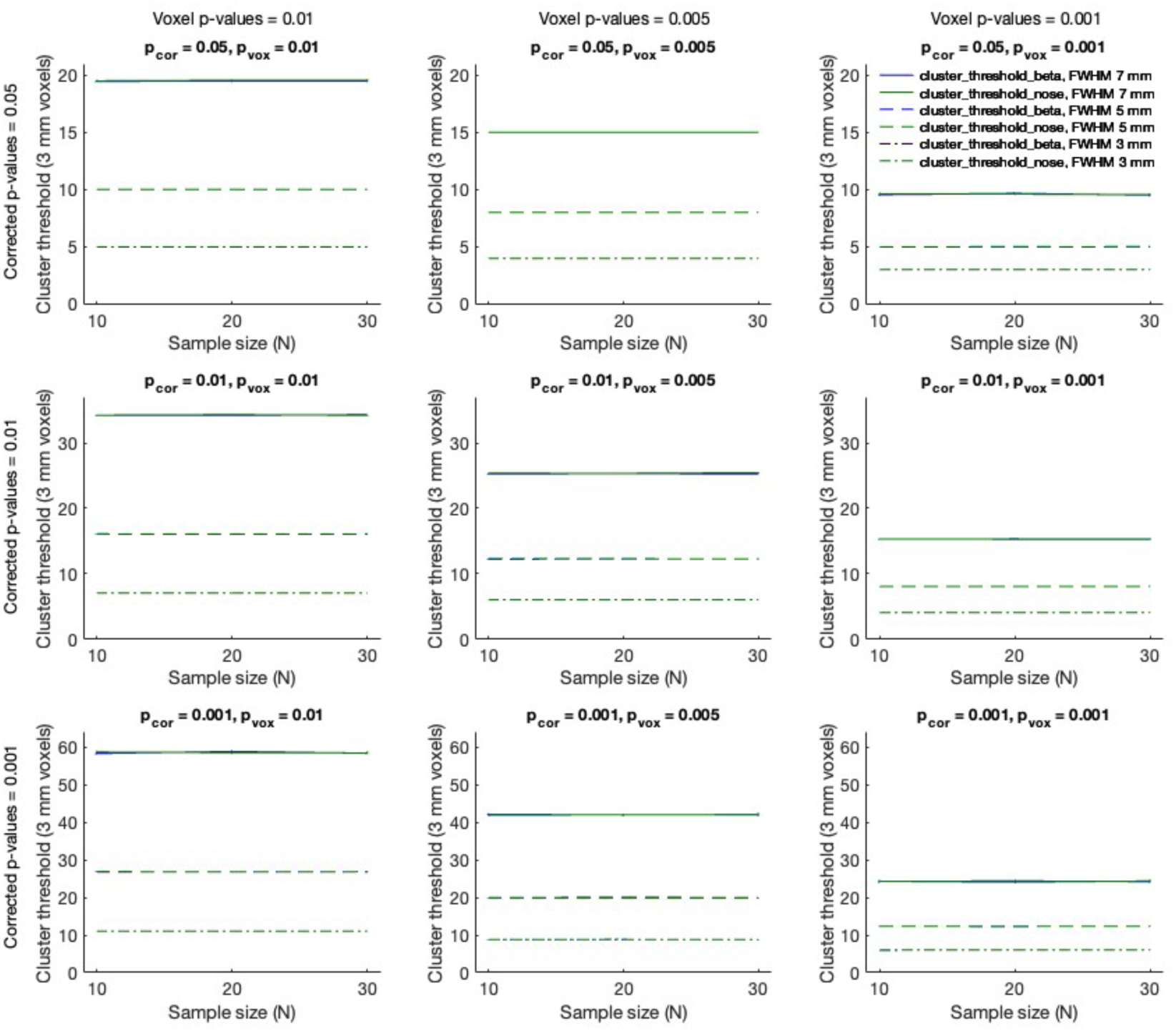
Cluster-extent thresholds (3 mm isotropic voxels) produced by cluster_threshold_beta (in blue) and cluster_threshold_nose (in green) as a function of corrected p-value (p_cor) and individual-voxel p-value (p_vox) with a full-width-half-maximum (FWHM) smoothing kernel of 3 mm, 5 mm, or 7 mm (key at the top right). In each panel, for each simulation set (N = 10, 20, and 30), the mean and 95% confidence intervals are shown, with lines connecting the means for each condition to illustrate trends as a function of N.

As illustrated in Figure 3, there did appear to be greater differential condition effects as a function of corrected p-values or individual-voxel p-values, which was expected. For corrected p-values, there was a significant main effect of corrected p-value (F(2,16194) = 2271.193, p < .001, η^2^ = .219) but no significant condition x corrected p-value interaction (F(2,16194) = .003, p = .997, η^2^ = 2.759x10^-7^). Bayesian analysis supported these findings as the corrected p-value model had a posterior model probability close to unity (P(M) = .983) and all other models had Bayes factor values close to zero (null model BF_10_ = 0.000, all other BF_10_ values < 0.018), which is very strong against the null model. For individual-voxel p-values, there was a significant main effect of individual-voxel p-value (F(2,16194) = 1187.255, p < .001, η^2^ = .128) but no significant condition x individual-voxel p-value interaction (F(2,16194) = 8.177x10^-4^, p =.999, η^2^ = 8.807x10^-8^). Bayesian analysis supported these findings as the individual-voxel p-value model had a posterior model probability close to unity (P(M) = .983) and all other models had Bayes factor values close to zero (null model BF_10_ = 0.000, all other BF_10_ values < 0.018), which is very strong evidence against the null model.

Figure 3 also appeared to show the expected greater differential condition effects with increasing FWHM smoothing kernels (which correspond to larger cluster thresholds). As a function of FWHM, there was a significant main effect of FWHM (F(2,16194) = 7052.289, p < .001, η^2^ = .466) but no significant condition x FWHM interaction (F(2,16194) = .002, p = .998, η^2^ = 1.259x10^-7^). Bayesian analysis supported these findings as the FWHM model had a posterior model probability close to unity (P(M) = .982) and all other models had Bayes factor values close to zero (null model BF_10_ = 0.000, all other BF_10_ values < .019), which is very strong evidence against the null model.

To quantify the similarity in cluster sizes in both conditions, for each simulation set, the cluster threshold produced by cluster_threshold_nose was divided by the cluster threshold produced by cluster_threshold_beta to produce a ratio. These results are illustrated in Supplemental Figure 1. This ratio ranged from 0.9904 to 1.0106, had a mean of 1.0008, and had a median of exactly 1. Thus, the average cluster threshold size produced by cluster_threshold_nose was nearly identical to the average cluster threshold size produced by cluster_threshold_beta.

## Simulation Set 3

The previous simulations were conducted using an acquisition volume with 3 mm isotropic voxels. With improvements in MRI scanner strength and fMRI acquisition protocols, smaller voxel sizes are becoming more common. In an effort to generalize the findings, this simulation set was conducted using the identical procedures as Simulation Sets 1 and 2, but the acquisition volume consisted of 2 mm isotropic voxels. As with the previous simulations, the hypotheses were that cluster_threshold_gamma would produce smaller cluster-extent thresholds than cluster_threshold_beta and that cluster_threshold_nose would produce the same cluster-extent thresholds as cluster_threshold_beta.

### Methods

Unless otherwise specified, the methods were identical to Simulation Sets 1 and 2, with the key difference being the acquisition volume voxel dimensions were set to 2 mm. To ensure similar whole-brain coverage to the previous simulations, matrix size was increased by a factor of 3/2 (rounded up) in all dimensions to 96 x 96 x 68 (i.e., 192 x 192 x 136 mm^3^, which has the identical spatial extent in the x- and y-dimensions and was 0.74% larger in the z-dimension than the previous simulations). The three conditions (cluster_threshold_beta, cluster_threshold_gamma, cluster_threshold_nose) required a total of 2,430,000 iterations with 33,210,000 simulated subjects.

### Results

Figure 4 shows the cluster-extent thresholds for each condition with FWHM smoothing kernel values of 3 mm, 5 mm or 7 mm. Of direct relevance to the primary hypothesis under investigation, there was a clear difference between conditions, with cluster_threshold_gamma (in red) producing smaller thresholds than both cluster_threshold_beta (in blue) and cluster_threshold_nose (in green; note, cluster threshold_beta and cluster_threshold_nose values were almost identical such that the blue lines were usually occluded by the green lines). Mirroring the order of analyses in the previous simulations, the statistical analysis first compared the cluster_threshold_beta and cluster_threshold_gamma conditions (as in Simulation Set 1). A frequentist analysis confirmed that there was a significant main effect of condition (for cluster_threshold_beta vs. cluster_threshold_gamma, F(1,16194) = 183.924, p < .001, η^2^ = .011), a significant main effect of N (F(2,16194) = 18.887, p < .001, η^2^ = .002), and a significant condition x N interaction (F(2,16194) = 19.227, p < .001, η^2^ = .002). A Bayesian analysis supported these findings as the interaction model (with Condition + N + Condition x N) had a posterior model probability close to unity (P(M) = 1.000), and all other models, including the null model, had Bayes factor values close to zero (all BF_10_ values < 3.184x10^-6^), which is very strong evidence against the null model. Follow-up analyses showed that cluster size was modulated by N in the cluster_threshold_gamma condition (frequentist, F(2,8097) = 46.326, p < .001, η^2^ = .011; Bayesian, N model P(M) = 1.00, null model BF_10_ = 8.022x10^-18^, which is very strong evidence against the null model), but was not modulated by N in the cluster_threshold_beta condition (frequentist, F(2,8097) = .005, p = .995, η^2^ = 1.194x10^-6^; Bayesian, null model P(M) = .999, N model BF_10_ = 0.001, which is very strong evidence favoring the null model).

**Figure 4.**
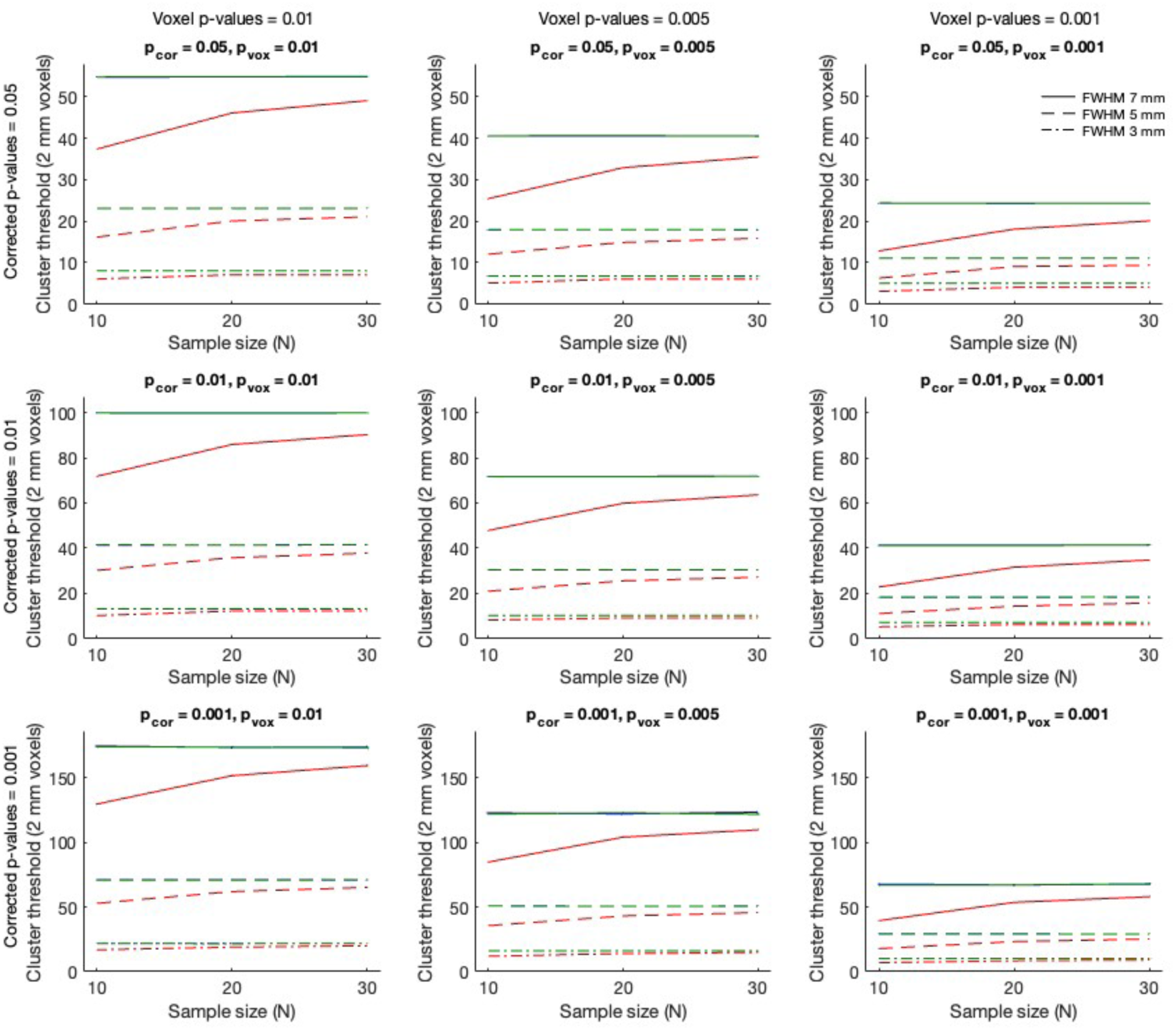
Cluster-extent thresholds (2 mm isotropic voxels) produced by cluster_threshold_beta (in blue), cluster_threshold_gamma (in red), and cluster_threshold_nose (in green) as a function of corrected p-value (p_cor) and individual-voxel p-value (p_vox) with a full-width-half-maximum (FWHM) smoothing kernel of 3 mm, 5 mm, or 7 mm (key at the top right). In each panel, for each simulation set (N = 10, 20, and 30), the mean and 95% confidence intervals are shown, with lines connecting the means for each condition to illustrate trends as a function of N.

As illustrated in Figure 4, there also appeared to be greater differential condition effects (between cluster_threshold_beta and cluster_threshold_gamma) for stricter corrected p-values, stricter individual-voxel p-values, and increasing FWHM smoothing kernels, which were expected. For corrected p-values, there was a significant main effect of corrected p-value (F(2,16194) = 1839.888, p < .001, η^2^ = .183) and a significant condition x corrected p-value interaction (F(2,16194) = 14.558, p < .001, η^2^ = .001; this is illustrated by parametrically increasing condition effects from top to bottom in the panels of Figure 4). Bayesian analysis supported these findings as the interaction model (with condition + corrected p-value + condition x corrected p-value) had a posterior model probability close to unity (P(M) = 1.000) and all other models had Bayes factor values close to zero (all BF_10_ values < 3.354x10^-4^), which is very strong evidence against the null model. For individual-voxel p-values, there was a significant main effect of individual-voxel p-value (F(2,16194) = 1116.789, p < .001, η^2^ = .120) but the condition x individual-voxel p-value interaction was not significant (F(2,16194) = 2.743, p = .064, η^2^ = 2.942x10^-4^; this is illustrated by the relatively similar condition effects from left to right in the panels of Figure 4). Bayesian analysis supported these findings as the condition + individual-voxel p-value model had a very high posterior model probability (P(M) = .978), the interaction model (with condition + individual-voxel p-value + condition x individual-voxel p-value) had the second highest posterior model probability (P(M) = .022), the sum of these posterior probabilities was close to unity (P(M) = 1.000), and all other models had Bayes factor values close to zero (all BF_10_ values < 7.145x10^-44^), which is very strong evidence against the null model. As a function of FWHM, there was a significant main effect of FWHM (F(2,16194) = 7171.198, p < .001, η^2^ = .462), and a significant condition x FWHM interaction (F(2,16194) = 93.317, p < .001, η^2^ = .006). Bayesian analysis supported these findings as the interaction model (with condition + FWHM + condition x FWHM) had a posterior model probability close to unity (P(M) = 1.000) and all other models had Bayes factor values close to zero (all BF_10_ values < 5.701x10^-38^), which is very strong evidence against the null model.

For this set of simulations, cluster thresholds produced by cluster_threshold_gamma were always smaller than those produced by cluster_threshold_beta. To quantify this effect, the cluster threshold produced by cluster_threshold_gamma was divided by the cluster threshold produced by cluster_threshold_beta to produce a ratio. These results are illustrated in Figures 5 for 3, 5, and 7 mm FWHM smoothing kernels. This ratio ranged from 0.5264 to 0.9291, had a mean of 0.8024, and had a median of 0.8392. Thus, the average cluster threshold size produced by cluster_threshold_gamma was 19.76% smaller than the average cluster threshold size produced by cluster_threshold_beta.

**Figure 5.**
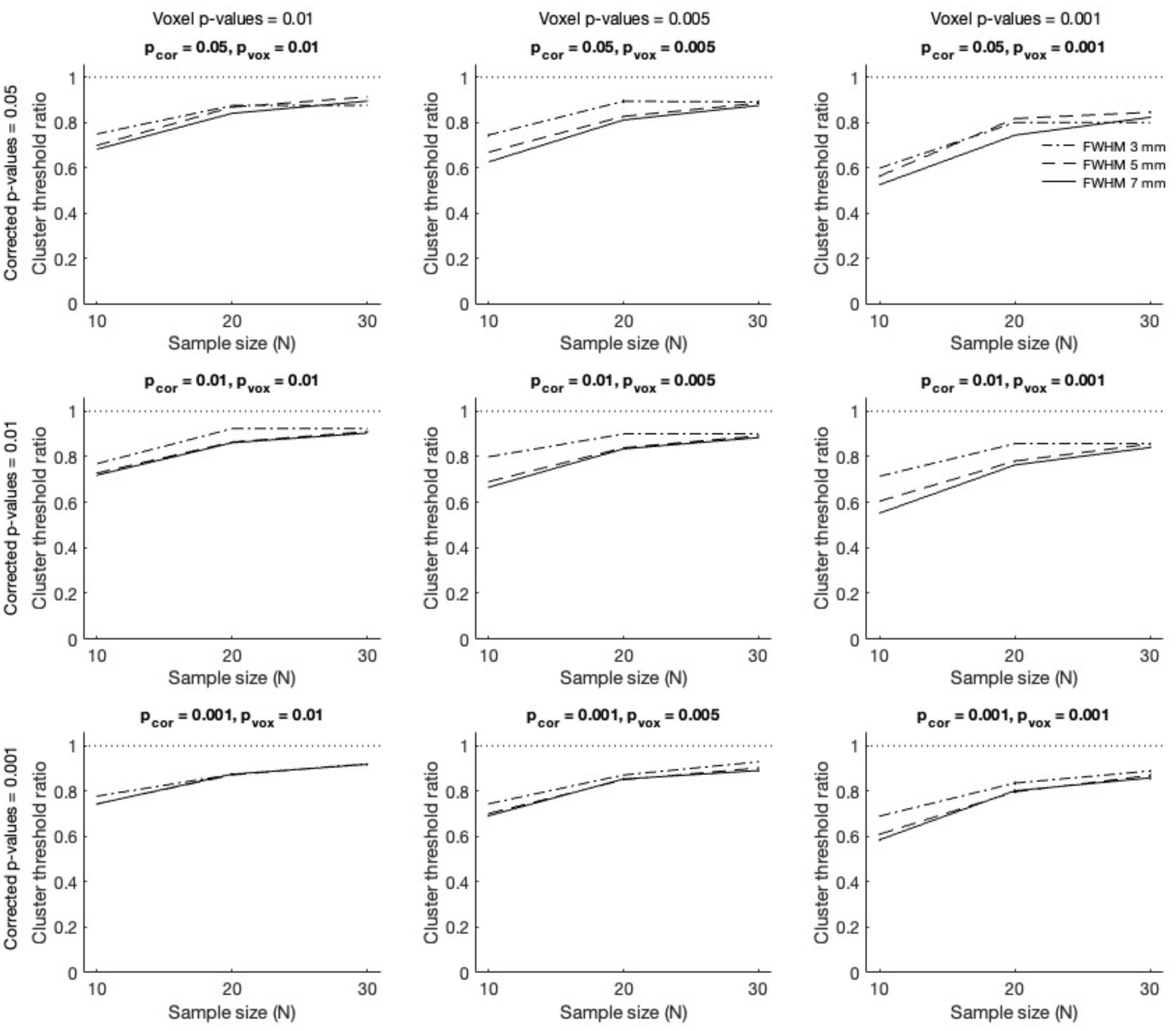
Ratio of cluster thresholds (2 mm isotropic voxels) produced by cluster_threshold_gamma and cluster_threshold_beta (solid line) as a function of corrected p-value (p_cor) and individual-voxel p-values (p_vox) with a FWHM smoothing kernel of 3 mm, 5 mm, or 7 mm (key at the top right; the dotted line demarcates a ratio of 1, for reference). In each panel, for each simulation set (N = 10, 20, and 30), the ratio of means and 95% confidence intervals are shown, with lines connecting the means for each condition to illustrate trends as a function of N.

Next, cluster_threshold_beta and cluster_threshold_nose were compared (as in Simulation Set 2). As illustrated in Figure 4, There was no clear difference between conditions, with cluster_threshold_nose (in green) producing identical or nearly identical thresholds as cluster_threshold_beta (in blue) for all comparisons (note, green lines with no hint of blue indicate perfect overlap). A frequentist analysis confirmed that there was no significant main effect of condition (cluster_threshold_beta vs. cluster_threshold_nose, F(1,16194) = .004, p =.951, η^2^ = 2.298x10^-7^), no significant main effect of N (F(2,16194) = .002, p = .998, η^2^ = 2.920x10^-7^), and no significant condition x N interaction (F(2,16194) = .003, p = .997, η^2^ = 3.234x10^-7^). A Bayesian analysis supported these findings as the null model had a posterior model probability close to unity (P(M) = .982) and all other models had Bayes factor values close to zero (all BF_10_ values < 0.019), which is strong evidence favoring the null model.

As expected, there appeared to be greater differential condition effects for stricter corrected p-values, stricter individual-voxel p-values, and increasing FWHM smoothing kernels. For corrected p-values, there was a significant main effect of corrected p-value (F(2,16194) = 1859.538, p < .001, η^2^ = .187) but no significant condition x corrected p-value interaction (F(2,16194) = .007, p = .993, η^2^ = 7.188x10^-7^). Bayesian analysis supported these findings as the corrected p-value model had a posterior model probability close to unity (P(M) = .983) and all other models had Bayes factor values close to zero (null model BF_10_ = 0.000, all other BF_10_ values < 0.019), which is very strong against the null model. For individual-voxel p-values, there was a significant main effect of individual-voxel p-value (F(2,16194) = 1042.773, p < .001, η^2^ = .114) but no significant condition x corrected p-value interaction (F(2,16194) = 5.436x10^-4^, p =.999, η^2^ = 5.948x10^-8^). Bayesian analysis supported these findings as the individual-voxel p-value model had a posterior model probability close to unity (P(M) = .983) and all other models had Bayes factor values close to zero (null model BF_10_ = 0.000, all other BF_10_ values < 0.018), which is very strong evidence against the null model. As a function of FWHM, there was a significant main effect of FWHM (F(2,16194) = 7928.393, p < .001, η^2^ = .495) but no significant condition x corrected p-value interaction (F(2,16194) = .004, p = .996, η^2^ = 2.68x10^-7^). Bayesian analysis supported these findings as the FWHM model had a posterior model probability close to unity (P(M) = .983) and all other models had Bayes factor values close to zero (null model BF_10_ = 0.000, all other BF_10_ values < .019), which is very strong evidence against the null model.

To quantify the similarity in cluster sizes in both conditions, for each simulation set, the cluster threshold produced by cluster_threshold_nose was divided by the cluster threshold produced by cluster_threshold_beta to produce a ratio. These results are illustrated in Supplemental Figure 2. This ratio ranged from 0.9885 to 1.0077, had a mean of 1.0001, and had a median of exactly 1. Thus, the average cluster threshold size produced by cluster_threshold_nose was nearly identical to the average cluster threshold size produced by cluster_threshold_beta.

## fMRI Analysis

The previous simulations provided support for the primary hypothesis that cluster_threshold_gamma would produce smaller cluster-extent thresholds than cluster_threshold_beta and cluster_threshold_nose and provided support for the secondary hypothesis that, for cluster_threshold_gamma, cluster-extent thresholds would increase as a function of sample size. The aim of this follow-up fMRI analysis was to confirm that the same pattern of results would be observed using a real fMRI dataset. Specifically, cluster thresholds produced by cluster_threshold_gamma and cluster_threshold_nose were directly compared. As mentioned previously, these scripts are nearly identical with the only difference being that, in cluster_threshold_nose, the between-subject mean value is not divided by the standard error term (i.e., voxel activity is simulated by the between-subject mean value rather than the corresponding t-value).

### Methods

Unless otherwise specified, the methods were identical to the previous simulations, employing slightly modified cluster_threshold_gamma and cluster_threshold_nose scripts, with the only difference being that the individual-subject brain activation volumes consisted of real fMRI data rather than simulated data.

Brain activation volumes from 54 participants were taken from one of our previously published spatial memory fMRI studies (Spets et al., 2019). Given the current aim, the methods will focus primarily on the fMRI dataset (for additional details, see Spets et al., 2019). During the encoding phase of each run, abstract shapes were presented to the left or right of fixation. During the corresponding retrieval phase of each run, previously presented abstract shapes were presented at fixation and, for each shape, participants made a “left”–“right” (spatial memory) classification. Functional and anatomic images were acquired on a Siemens 3-Tesla scanner with a head coil using a T2*-weighted echo-planar imaging sequence (TR = 2000 ms, TE = 30 ms, 64 x 64 acquisition matrix, 26–30 slices, 4.5 mm isotropic resolution). Analysis was conducted using BrainVoyager (Maastricht, The Netherlands). Images were slice-time corrected, motion corrected, temporal components below 2 cycles per run length were removed, voxels were resampled at 3 mm^3^, and volumes were transformed into Talairach space (to maximize spatial resolution, spatial smoothing was not conducted). For each participant, a general linear model analysis was conducted by modeling responses to all event types. Of relevance to the current analysis, the contrast of accurate spatial memory responses during retrieval (i.e., responding “left” for items previously presented on the left, “left”/left, or “right”/right) and inaccurate spatial memory responses during retrieval (i.e., “left”/right or “right”/left) was used to produce a spatial memory hit vs. miss brain activation volume for each participant. Brain activation volumes from 36 females and 18 males were included in the analysis (these were the same participants included in the results of Spets et al., 2019).

The cluster_threshold_gamma and cluster_threshold_nose simulations were identical to that described in Simulation Set 1 and Simulation Set 2 except that, for each participant, a brain activation volume was read into the script rather than simulating a brain activation volume. For each iteration, a random set of participants was selected with the constraint that, for each sample size (10, 20, or 30), half of the participants were female, and half of the participants were male (to avoid oversampling the larger set of females). Although the same pool of participants was sampled for each iteration, such Bootstrap procedures can be used to provide an estimate of variance (i.e., see Efron & Tibshirani, 1993). As with the simulations, for each corrected p-value and individual-voxel p-value, 100 iterations were repeated 100 times. Given that real fMRI data has inherent spatial autocorrelation, additional convolution with Gaussian smoothing kernels was not conducted (i.e., the FWHM parameter was set to zero in the scripts). As this was a follow-up analysis based on the complete set of simulations, to minimize the number of comparisons, in an effort to minimized type I error, statistical tests were limited to those directly related to the two hypotheses under investigation.

### Results

Figure 6 shows the cluster-extent thresholds for each condition. Of direct relevance to the primary hypothesis under investigation, there was a clear difference between conditions, with cluster_threshold_gamma (in red) producing smaller thresholds than cluster_threshold_nose (in green) for all of the comparisons. A frequentist analysis confirmed that there was a significant main effect of condition (cluster_threshold_nose vs. cluster_threshold_gamma, F(1,5398) = 411.207, p < .001, η^2^ = .071). A Bayesian analysis supported this finding as the Condition model had a posterior model probability close to unity (P(M) = 1.000), and the null model had a Bayes factor value close to zero (BF_10_ = 4.449x10^-85^), which is very strong evidence against the null model. In support of the secondary hypothesis, cluster size was modulated by N in the cluster_threshold_gamma condition (frequentist, F(2,2697) = 54.939, p < .001, 1^2^ = .039; Bayesian, N model P(M) = 1.00, null model BF_10_ = 1.615x10^-21^, which is very strong evidence against the null model).

**Figure 6.**
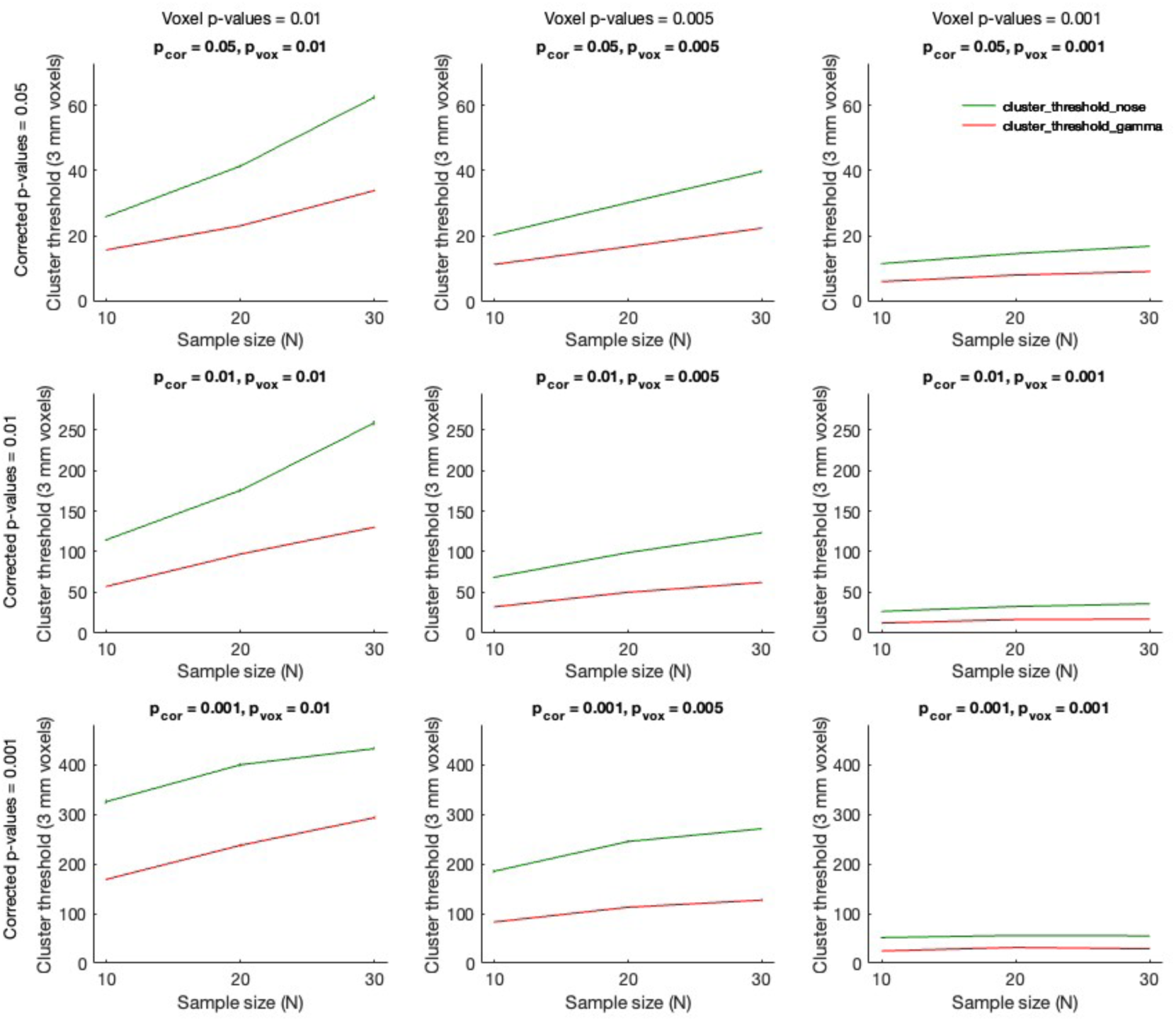
Cluster-extent thresholds (3 mm isotropic voxels) produced by cluster_threshold_nose (in green) and cluster_threshold_gamma (in red) as a function of corrected p-value (p_cor) and individual-voxel p-value (p_vox; key at the top right). In each panel, for each fMRI data set (N = 10, 20, and 30), the mean and 95% confidence intervals are shown, with lines connecting the means for each condition to illustrate trends as a function of N.

It is notable that cluster_threshold_nose values also appeared to increase as a function of sample size. This can be attributed to consistent activity (associated with spatial memory hits vs. misses) across participants, where averaging over a higher number of participants would be expected to produce larger regions of contiguous activity (yielding larger cluster-extent thresholds). However, this effect reflects employing brain activation volumes that were spatially correlated, rather than null data. While this is an important topic of future research (see the Discussion), it is not of relevance to the two hypotheses under investigation.

Cluster thresholds produced by cluster_threshold_gamma were always smaller than those produced by cluster_threshold_nose. To quantify this effect, the cluster threshold produced by cluster_threshold_gamma was divided by the cluster threshold produced by cluster_threshold_nose to produce a ratio. These results are illustrated in Figure 7. This ratio ranged from 0.4498 to 0.6784, had a mean of 0.5262, and had a median of 0.5213. Thus, the average cluster threshold size produced by cluster_threshold_gamma was 47.38% smaller than the average cluster threshold size produced by cluster_threshold_nose.

**Figure 7.**
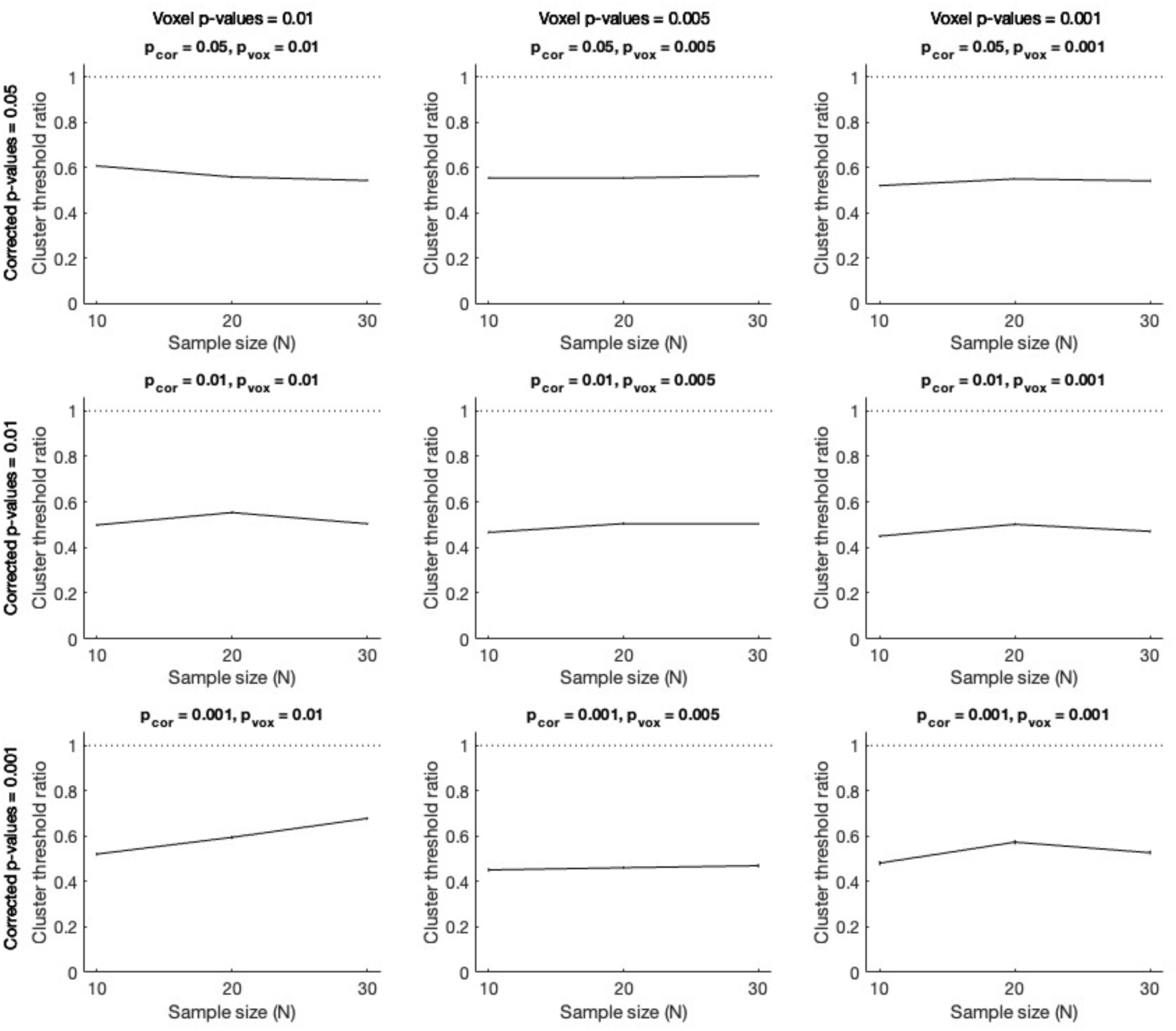
Ratio of cluster thresholds (3 mm isotropic voxels) produced by cluster_threshold_gamma and cluster_threshold_nose as a function of corrected p-value (p_cor) and individual-voxel p-value (p_vox; key at the top right; the dotted line demarcates a ratio of 1, for reference). In each panel, for each fMRI data set (N = 10, 20, and 30), the ratio of means and 95% confidence intervals are shown, with lines connecting the means for each condition to illustrate trends as a function of N.

## Discussion

The primary hypothesis of the present investigation was that incorporating sample size into the simulations would reduce the cluster-extent threshold. It was further hypothesized that this reduction in cluster-extent threshold was due to between-subject variability. The identical pattern of results was obtained in Simulation Sets 1 and 2, which used an isotropic voxel size of 3 mm, and Simulation Set 3, which used an isotropic voxel size of 2 mm, such that the results of all simulations will be discussed together. In support of the first hypothesis (tested in Simulation Sets 1 and 3), frequentist analyses showed a significant effect of condition (cluster_threshold_beta vs. cluster_threshold_gamma). Specifically, cluster_threshold_gamma clusters were about 18% smaller than cluster_threshold_beta clusters. There was also a significant effect of sample size, showing that cluster-extent threshold was affected by the specific sample size, and a significant condition x sample size interaction. The interaction was driven by significant increases in cluster_threshold_gamma thresholds for larger sample sizes, which can be attributed to lower standard error values resulting in higher t-values (and thus more voxels being classified as significant). All the preceding findings were supported by Bayesian analyses. To test the second hypothesis, between-subject variability modeled in cluster_threshold_gamma was eliminated by removing the standard error term to create cluster_threshold_nose (in Simulation Sets 2 and 3). In support of this hypothesis, frequentist analyses showed no significant effects of condition (cluster_threshold_beta vs. cluster_threshold_nose), sample size, or their interaction. Of importance, the Bayesian analyses produced very strong evidence favoring the null model. These simulation results were replicated in a real fMRI data set. These findings show that sample size is an important factor in fMRI methods that correct for multiple comparisons and provides an explanation of how sample size modulates cluster-extent threshold.

In addition to the main hypotheses under investigation, the effects of condition and corrected p-value, individual-voxel p-value, and FWHM were evaluated. As expected, stricter corrected p-values were associated with larger cluster-extent thresholds, stricter individual-voxel p-values were associated with smaller cluster-extent thresholds, and larger FWHM values were associated with larger cluster-extent thresholds. For the cluster_threshold_beta versus cluster_threshold_gamma comparison, there were significant condition x corrected p-value and condition x FWHM interactions, but the condition x individual-voxel p-value interaction was not significant. For the cluster_threshold_beta versus cluster_threshold_nose comparison, none of these interactions were significant. Bayesian analyses supported these findings. These results show that sample size is a critical parameter that can interact with other parameters. Thus, incorporating sample size into algorithms that correct for multiple comparisons in fMRI analysis is necessary to accurately estimate cluster-extent threshold. Toward this aim, cluster_threshold_gamma is now available for use by the neuroimaging community (Slotnick, 2025b).

### Other methods should incorporate sample size

More broadly, the present finding that sample size can affect cluster-extent threshold should be incorporated into other methods to reduce type II error. This includes Monte Carlo simulation methods such as the original version of AFNI (Cox, 1996) and the currently recommended ‘long-tail’ version of AFNI (Cox et al., 2017). For nonparametric permutation methods (e.g., Eklund et al., 2017), the actual number of participants corresponding to the to-be-corrected study should be employed, rather than an arbitrary sample size that assumes the value of this parameter will not affect the results. If sample size is not implemented in these methods, the present findings indicate that this may result in increased type II error due to overly strict thresholds (e.g., in the case of AFNI, larger than optimal cluster thresholds).

### Assuming non-task activity reflects null data is questionable

Analysis methods that employ the group residual activation map, including SPM (Kiebel et al., 1999) and FSL (Jenkinson et al., 2012), implicitly take sample size into account. However, these methods assume that the residual map reflects null data, which is questionable given that such data is defined as non-task activity and thus may reflect activity in the default-mode network (Slotnick, 2017a, 2017b). It is important to highlight that in some experimental designs, the residual activation map may reflect null data, while in other experimental designs, the residual map may reflect robust default-mode network activity. This likely depends on the degree to which participants have time between trials to engage in non-task processing. Critically, given the wide range of experimental designs in the field, the residual activation map is expected to reflect default-network activity in many studies. Determining how often to what degree this occurs is an important topic for future research.

The default network has been associated with many cognitive processes such as planning for the future, daydreaming, mind wandering, lapses of attention, and retrieval of personal information (for a review, see Buckner, Andrews-Hanna, & Schacter, 2008). As illustrated in Figure 8a, non-task/resting-state activity is associated with robust activity in many brain regions including the dorsolateral prefrontal cortex, the medial prefrontal cortex, the inferior parietal cortex, and the medial parietal cortex. Methods that assume non-task data, such as the group residual map in SPM and FSL, reflects null activity, along with those that explicitly assume resting-state data reflects null activity (Eklund et al., 2017, 2019) or model this type of brain activity (i.e., the long-tail version of AFNI; Cox et al., 2017), may be enforcing stricter-than-optimal thresholds resulting in increased type II error. Figure 8, panels b and c, illustrate “false clusters” from two different datasets (Eklund et al., 2019) that clearly reflect default-network activity (compare Figures 8b and 8c to Figure 8a). Although these default-network activations are sometimes referred to as “false clusters” (Eklund et al., 2017, 2019), they actually reflect true clusters that will artifactually inflate cluster-extent thresholds (Slotnick, 2017a; the primary one-tailed t-tests results in Eklund et al., 2017, also appeared to be artifactually inflated; Flandin & Friston, 2019; Slotnick, 2017b).

**Figure 8.**
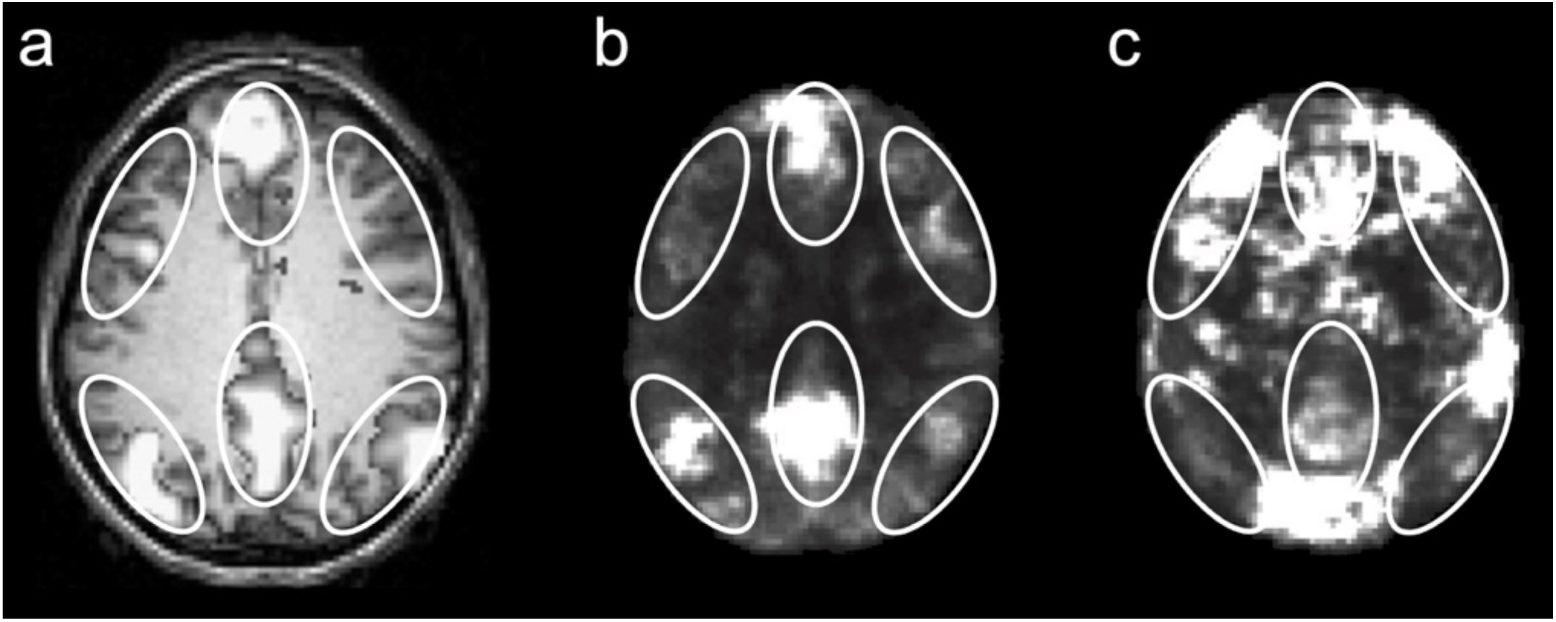
fMRI default-network activity and resting-state activity that has been assumed to reflect null data. (a) Default-network activity (ovals demarcate default-network regions; axial view, anterior toward the top; adapted from Figure 3 in NeuroImage, 37/4, Randy L. Buckner and Justin L. Vincent, Unrest at rest: Default activity and spontaneous network correlations, 1091–1096, 2007, with permission from Elsevier). (b) Resting-state “false clusters” from the Cambridge dataset (ovals copied from panel a). (c) Resting state “false clusters” from the Oulu dataset (ovals copied from panel a; b and c were adapted from Eklund et al., 2019, Figure 7 and Figure 8, respectively; Creative Commons CC BY license).

As highlighted in Slotnick (2017a, 2017b), a more accurate way to model null data would be to contrast odd versus even trials of one event type. Ideally, this event should correspond to an isolated cognitive process, with minimal true brain activations, to avoid variability between regressors. Resting-state (or residual) activation maps would be a poor choice given that default-network activity has a broad spatial extent, high magnitude, and high variability (Mao et al., 2015; Figure 8c shows an extreme example). After contrasting well-matched event types (corresponding to minimal activity), it would be expected that true null data would have a relatively uniform distribution across the brain (as it does using the methods employed in the current simulations; see Supplemental Figure 3).

It should be noted that the newest method of the AFNI program (2025b), although not recommended at this time, takes sample size into account and makes a laudable attempt to create a true null volume. Specifically, the procedure generates the null volume by randomizing the sign of the residuals for each subject (Cox et al., 2017). Assuming that residuals reflect a stable pattern of activity in particular regions (that may or may not reflect default-network activity, as discussed above), this procedure should yield approximately half of the subject with increases in activity in these regions, and the other half of the subjects with decreases in activity in these regions, the sum of which should produce activations that vary around zero in these and other regions. As mentioned above, activity in the default-network regions can be highly variable and thus, if residuals do reflect default-network activity, this procedure may produce significant differences and inflate cluster-extent thresholds. Still, this is the only method employing residuals that may actually eliminate true activity from the analysis and thus is arguably the most accurate of such methods available. To avoid the potential issues with default-network activity described above, it would likely be more accurate to conduct a similar procedure but use the single event regressor (of all regressors in a particular study) that produces the least number of significant activations (i.e., is most similar to null a priori) and assign exactly half of the participants with a positive sign and the other half with a negative sign, the sum of which should create the most accurate null volume for that dataset. Optimizing procedures that estimate the null volume to correct for multiple comparisons is an important topic for future research.

### More degrees of freedom, more problems

For a given experiment, type I error and type II error are linked. At the desired individual-voxel p-value, if the corrected p-value is made stricter, type I error will decrease and type II error will increase, while if the corrected p-value is made more lenient, type I error will increase and type II error will decrease. Although there is a general focus in the field on reducing type I error, it is now required to correct for multiple comparisons in an acceptable manner for an fMRI study to be published in a respectable journal. That is, at this time, familywise type I error is required to be controlled in virtually all fMRI studies. As such, type II error should be considered a more pressing problem in the field.

One way to reduce type II error is to increase sample size. For a given study, the dual aims of generalizing to the population and increasing power (i.e., detecting an effect when it exists; 1 – type II error) tend to boost sample size, while the high cost of conducting fMRI studies tends to restrict sample size. Button et al. (2013) estimated that the average statistical power of neuroscience studies, which included an analysis of brain volume abnormality studies and animal studies, was between 8 and 31% and to achieve a power of 80% in different rodent maze completion experiments would require 68-134 subjects (an analysis of fMRI data was not conducted). Thus, approximately 100 subjects were recommended to achieve sufficient power. Aligning with Button et al., Poldrack et al. (2017) showed that median sample size of fMRI studies has increased from 9 in 1996 to 28.5 in 2015 and estimated the corresponding effects sizes needed to achieve 80% power ranged from about 1.25 to 0.75 (i.e., only large effect sizes had sufficient power for true effects to be detected). Poldrack et al. argued that much larger sample sizes than are typically utilized are needed to achieve sufficient power, acknowledging “given that research funding will probably not increase to accommodate these larger samples, fewer studies may be funded, and researchers with fewer resources may have a more difficult time performing research that meets these standards” (p. 117). Both Button et al. and Poldrack et al. made many assumptions when computing sample size estimates to achieve 80% power and, at least for fMRI, the estimates seem at odds with reality. For example, in 2020, approximately 50% of all episodic memory studies reported activity in the hippocampus (Slotnick, 2023), even though fMRI studies almost never have large sample sizes due to the high cost off scanning. This suggests that fMRI studies with typical sample sizes (e.g., 20 participants) usually have sufficient power to detect activity of interest. Even if the power estimates of Button et al. and Poldrack et al. were correct, other factors need to be considered when deciding whether to conduct one large study versus several smaller studies.

Lieberman and Cunningham (2009) highlighted four negative consequences on focusing on only type I error including: 1) “increased type II errors”, 2) “a bias toward publishing large and obvious effects”, 3) “a bias against observing effects associated with complex cognitive and affective processes”, and 4) “deficient meta-analyses” (p. 424). Along the same lines, Hopfinger (2017) underscored that “replication across labs remains a most critical part of scientific study” and “an overemphasis on increasingly conservative thresholds can negatively impact the potential for replication of studies and the pursuit and reporting of innovative results” (p. 145). Said another way, if one large study is conducted from a single laboratory (with say, 100 participants) versus many studies from different laboratories (e.g., with 20 participants), this will stifle meta-analyses, replication, and innovation. Relating to points 1 and 2 of Lieberman and Cunningham, we recently published an fMRI study with 864 participants (mining data from the Human Connectome Project; Fritch et al., 2025). At a standard individual-voxel threshold of p < .001, a general linear model analysis produced activity across the entire brain, which can be attributed to the massive degrees of freedom. A p-value < 1 x 10^-30^ was required for the results to be interpretable, which necessarily translated in a high degree of type II error (these results were not included in Fritch et al., 2025). It should be noted that this potential problem of increased type II error in fMRI studies with large sample sizes does not apply to the simulation sets conducted in the current study. For simulations, running 10,000 iterations is the gold standard to produce results that converge on a stable solution, which, in this study, translated into millions of simulated subjects. However, rather than conducting a conventional fMRI general linear model analysis (with a single individual-voxel p-value and a single corrected p-value), the simulations parametrically varied key parameters and evaluated the stable patterns of results (and such stability requires a large number of simulated subjects).

A related issue is that large fMRI datasets available for mining (with real participants) cannot be used to evaluate specific hypotheses that are of interest to the large majority of scientists. Mining such data puts the answer (i.e., the data from someone else’s experiment) before the question, which is not an efficient way to conduct science, where an experiment should be intelligently designed to adjudicate between critical hypotheses (Platt, 1964). Considering the factors above, although at first blush it may seem best to conduct a lower number of fMRI studies with larger sample sizes, it seems prudent to conduct a higher number of fMRI studies with sufficient sample sizes to maximize the chances of revealing important effects in a timely manner. Determining optimal sample size is an important topic for future research.

With the understanding that there are constraints on fMRI sample sizes, Cunningham and Koscik (2017) proposed a novel method to reduce type II error. Given that certain regions are expected to be active in a given experiment, those regions can be subjected to a more lenient threshold (thus decreasing type II error), whereas the error in other regions (e.g., the rest of the brain) could be subjected to a stricter threshold such that the familywise type I error rate across the entire brain would be set to the desired level. This algorithm, or other to-be-derived algorithms that may reduce type II error while controlling familywise type I error, should be incorporated into widely available analysis software or scripts.

### Conclusion

The current study showed that increasing sample size can reduce the cluster-extent threshold. Employing these smaller cluster thresholds (that can be easily computed with the cluster_threshold_gamma script; Slotnick, 2025b) will reduce type II error rate, while still maintaining the desired type I error rate. The present finding has ramifications for methods that do not incorporate sample size into their computations, and this should be done to reduce type II error. It was also argued that fMRI group residuals and resting-state data may not reflect true null volumes, employing such volumes could increase type II error, and that null volumes should be estimated using appropriate techniques. Our overall aim should be to optimize the methods we employ to balance type I error with type II error. By doing so, the field can make important discoveries as rapidly as possible.

## Acknowledgments

I thank the editor and two anonymous reviewers for their insightful comments.

## Supplemental Figures

**Supplemental Figure 1.**
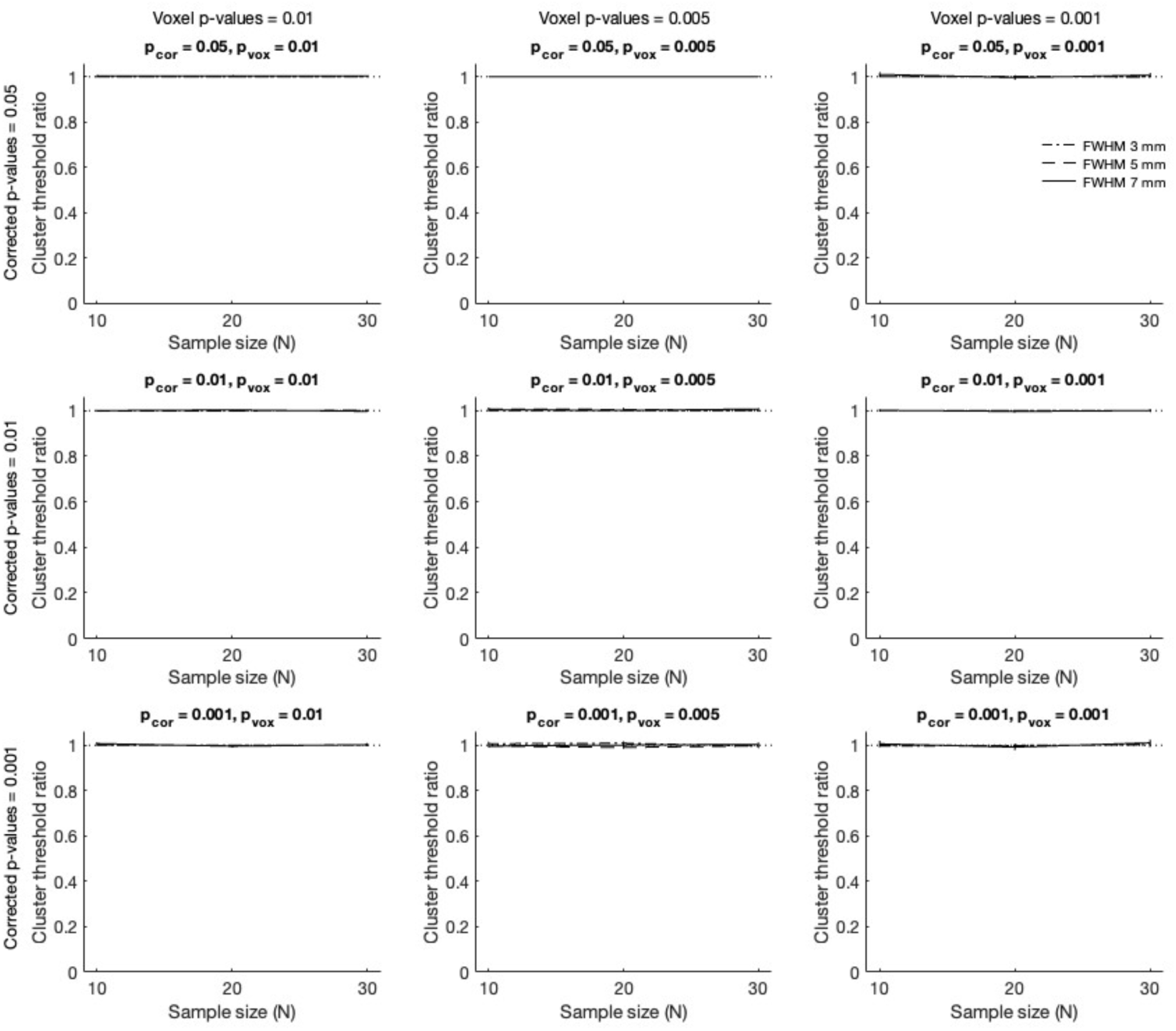
Ratio of cluster thresholds (3 mm isotropic voxels) produced by cluster_threshold_nose and cluster_threshold_beta as a function of corrected p-value (p_cor) and individual-voxel p-value (p_vox) with a FWHM smoothing kernel of 3 mm, 5 mm, or 7 mm (key at the top right; the dotted line demarcates a ratio of 1, for reference). In each panel, for each simulation set (N = 10, 20, and 30), the ratio of means and 95% confidence intervals are shown, with lines connecting the means for each condition to illustrate trends as a function of N.

**Supplemental Figure 2.**
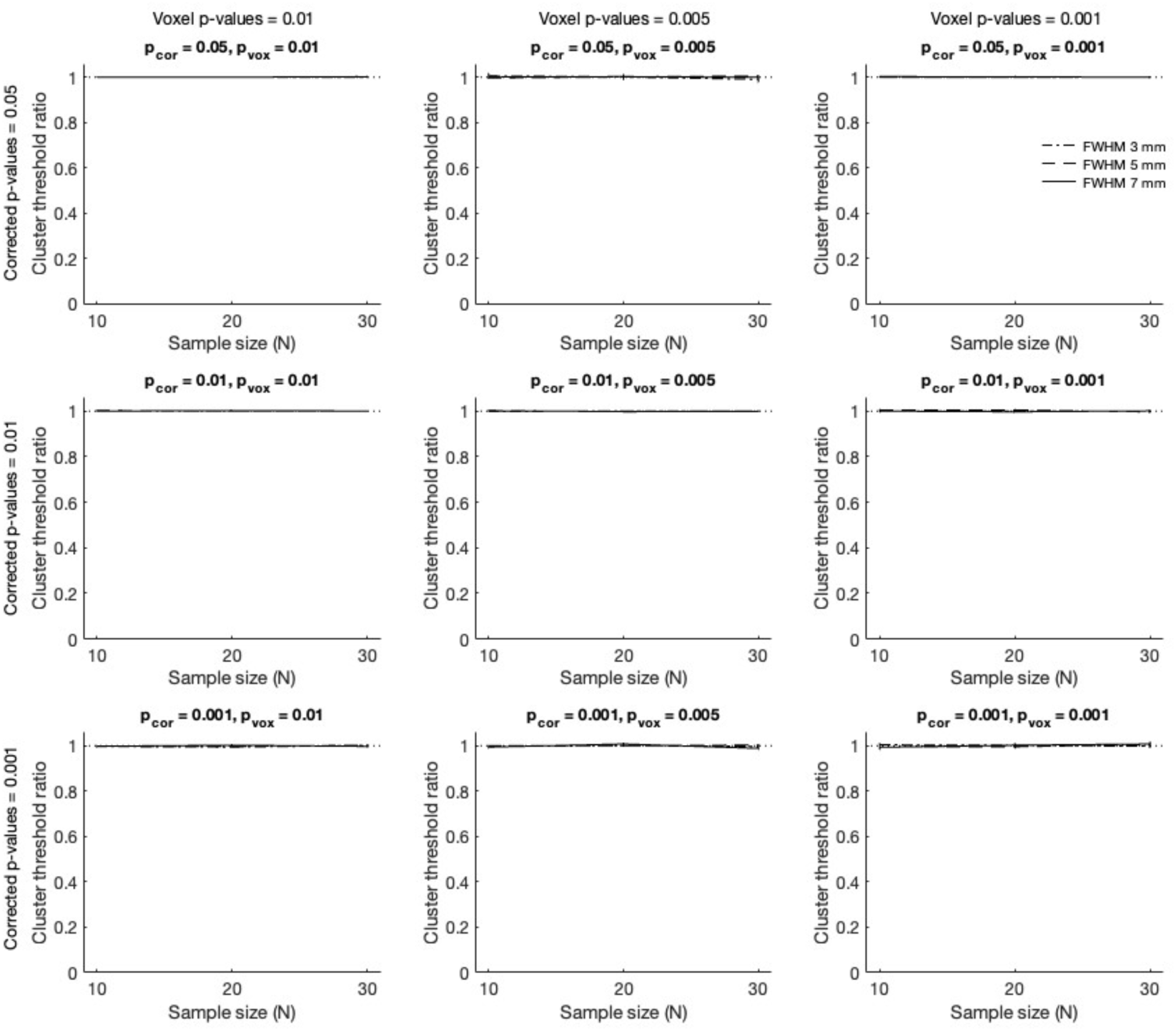
Ratio of cluster thresholds (2 mm isotropic voxels) produced by cluster_threshold_nose and cluster_threshold_beta as a function of corrected p-value (p_cor) and individual-voxel p-value (p_vox) with a FWHM smoothing kernel of 3 mm, 5 mm, or 7 mm (key at the top right; the dotted line demarcates a ratio of 1, for reference). In each panel, for each simulation set (N = 10, 20, and 30), the ratio of means and 95% confidence intervals are shown, with lines connecting the means for each condition to illustrate trends as a function of N.

**Supplemental Figure 3.**
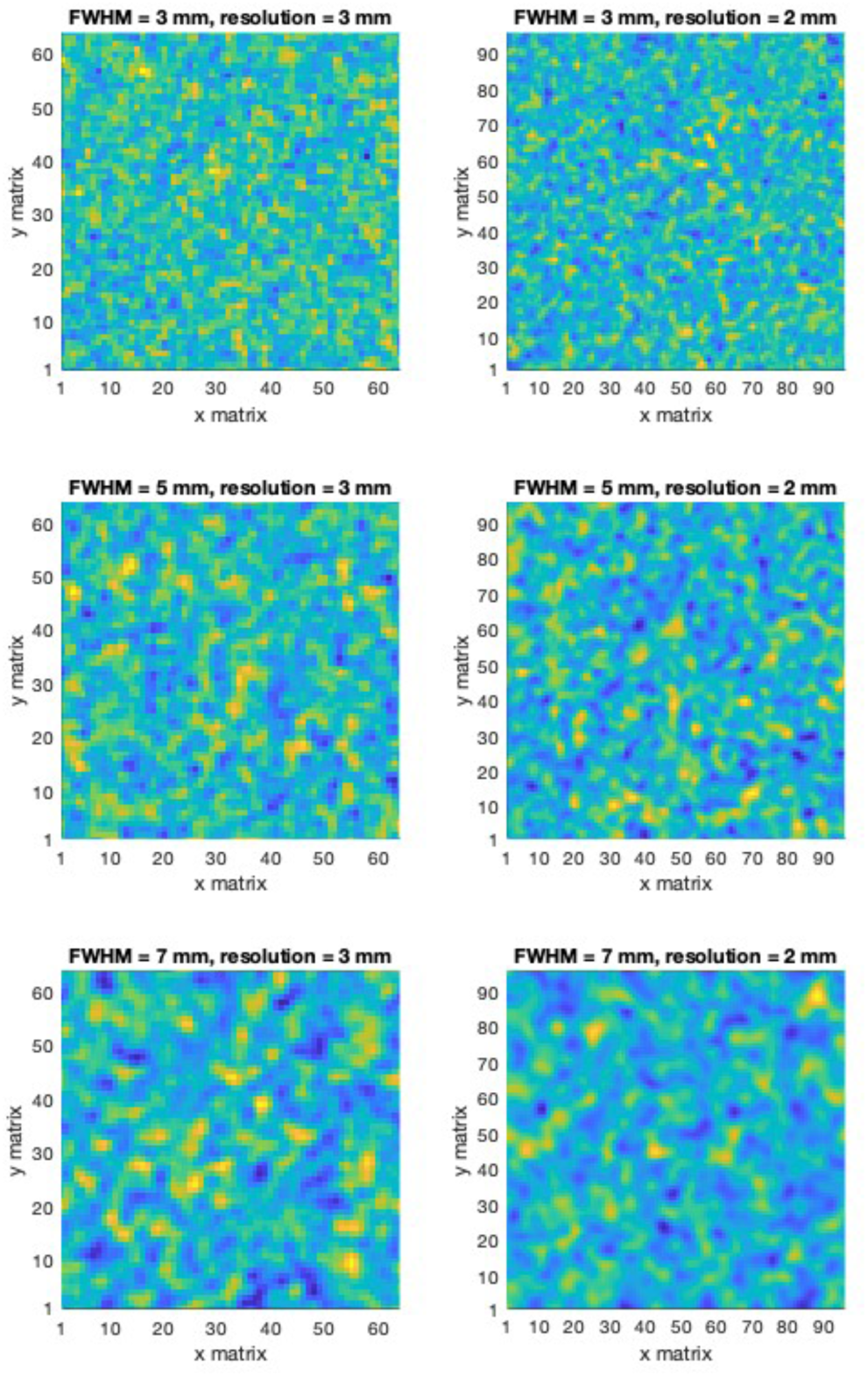
Mean amplitude of null volumes (20 subjects, middle slice) for 3 mm voxel resolution (left column) and 2 mm voxel resolution (right column) with smoothing kernels of 3, 5, and 7 mm (top, middle, and bottom rows). A random number was used to seed the random number generator and then each null volume was created, in a single iteration, using cluster_threshold_gamma as described in the Methods of Simulation Set 1 and Simulation Set 3 (images were not selected to identify “representative” results).

## References

AFNI program. (2025a, Jul 11). 3dClustSim. Retrieved from https://afni.nimh.nih.gov/pub/dist/doc/program_help/3dClustSim.html

AFNI program. (2025b, Jul 11). 3dttest++. Retrieved from https://afni.nimh.nih.gov/pub/dist/doc/program_help/3dttest++.html

Benjamini Y, Hochberg Y. Controlling the false discovery rate: a practical and powerful approach to multiple testing. J R Statist Soc B. 1995 Jan;57(1):289–300. doi: 10.1111/j.2517-6161.1995.tb02031.x

Benjamini Y, Krieger AM, Yekutieli D. Adaptive linear step-up procedures that control the false discovery rate. Biometrika. 2006 Sep;93(3):491–507. doi: 10.1093/biomet/93.3.491

Buckner RL, Andrews-Hanna JR, Schacter DL. The brain’s default network: anatomy, function, and relevance to disease. Ann N Y Acad Sci. 2008 Mar;1124:1–38. doi: 10.1196/annals.1440.011

Buckner RL, Vincent JL. Unrest at rest: default activity and spontaneous network correlations. Neuroimage. 2007 Oct 1;37(4):1091–6; discussion 1097-9. doi: 10.1016/j.neuroimage.2007.01.010

Button KS, Ioannidis JP, Mokrysz C, Nosek BA, Flint J, Robinson ES, Munafò MR. Power failure: why small sample size undermines the reliability of neuroscience. Nat Rev Neurosci. 2013 May;14(5):365–76. doi: 10.1038/nrn3475

Cox RW. AFNI: software for analysis and visualization of functional magnetic resonance neuroimages. Comput Biomed Res. 1996 Jun;29(3):162–73. doi: 10.1006/cbmr.1996.0014

Cox RW, Chen G, Glen DR, Reynolds RC, Taylor PA. FMRI Clustering in AFNI: False-Positive Rates Redux. Brain Connect. 2017 Apr;7(3):152–171. doi: 10.1089/brain.2016.0475

Cunningham WA, Koscik TR. Balancing Type I and Type II error concerns in fMRI through compartmentalized analysis. Cogn Neurosci. 2017 Jul;8(3):147–149. doi: 10.1080/17588928.2017.1299122

Efron B, Tibshirani RJ (1993). An introduction to the Bootstrap. Chapman and Hall.

Eklund A, Knutsson H, Nichols TE. Cluster failure revisited: Impact of first level design and physiological noise on cluster false positive rates. Hum Brain Mapp. 2019 May;40(7):2017–2032. doi: 10.1002/hbm.24350

Eklund A, Nichols TE, Knutsson H. Cluster failure: Why fMRI inferences for spatial extent have inflated false-positive rates. Proc Natl Acad Sci U S A. 2016 Jul 12;113(28):7900–5. doi: 10.1073/pnas.1602413113

Flandin G, Friston KJ. Analysis of family-wise error rates in statistical parametric mapping using random field theory. Hum Brain Mapp. 2019 May;40(7):2052–2054. doi: 10.1002/hbm.23839

Fritch HA, Steinkrauss AC, Slotnick SD. Resting-State Functional Connectivity With the Anterior and Posterior Hippocampus: An Analysis of fMRI Data From the Human Connectome Project. Hippocampus. 2025 Jul;35(4):e70023. doi: 10.1002/hipo.70023

Holmes DT, Buhr KA. Error propagation in calculated ratios. Clin Biochem. 2007 Jun;40(9-10):728–34. doi: 10.1016/j.clinbiochem.2006.12.014

Hopfinger JB. Replication and innovation versus a perfect ‘.05’. Cogn Neurosci. 2017 Jul;8(3):145–147. doi: 10.1080/17588928.2017.1297296

Hopfinger JB, Büchel C, Holmes AP, Friston KJ. A study of analysis parameters that influence the sensitivity of event-related fMRI analyses. Neuroimage. 2000 Apr;11(4):326–33. doi: 10.1006/nimg.2000.0549. PMID: 10725188.

JASP Team (2025). JASP (Version 0.19.3)[Computer software]

Jenkinson M, Beckmann CF, Behrens TE, Woolrich MW, Smith SM. FSL. Neuroimage. 2012 Aug 15;62(2):782-90. doi: 10.1016/j.neuroimage.2011.09.015

Kass RE, Raftery AE. Bayes factors. J Am Stat Assoc. 1995 Jun;90(430):773–795.

Kiebel SJ, Poline JB, Friston KJ, Holmes AP, Worsley KJ. Robust smoothness estimation in statistical parametric maps using standardized residuals from the general linear model. Neuroimage. 1999 Dec;10(6):756–66. doi: 10.1006/nimg.1999.0508

Lieberman MD, Cunningham WA. Type I and Type II error concerns in fMRI research: re-balancing the scale. Soc Cogn Affect Neurosci. 2009 Dec;4(4):423–8. doi: 10.1093/scan/nsp052

Mao D, Ding Z, Jia W, Liao W, Li X, Huang H, Yuan J, Zang YF, Zhang H. Low-Frequency Fluctuations of the Resting Brain: High Magnitude Does Not Equal High Reliability. PLoS One. 2015 Jun 8;10(6):e0128117. doi: 10.1371/journal.pone.0128117

Nichols TE, Eklund A, Knutsson H. A defense of using resting-state fMRI as null data for estimating false positive rates. Cogn Neurosci. 2017 Jul;8(3):144–149. doi: 10.1080/17588928.2017.1287069

Platt JR. Strong Inference: Certain systematic methods of scientific thinking may produce much more rapid progress than others. Science. 1964 Oct 16;146(3642):347–53. doi: 10.1126/science.146.3642.347

Poldrack RA, Baker CI, Durnez J, Gorgolewski KJ, Matthews PM, Munafò MR, Nichols TE, Poline JB, Vul E, Yarkoni T. Scanning the horizon: towards transparent and reproducible neuroimaging research. Nat Rev Neurosci. 2017 Feb;18(2):115–126. doi: 10.1038/nrn.2016.167

Slotnick SD. Resting-state fMRI data reflects default network activity rather than null data: A defense of commonly employed methods to correct for multiple comparisons. Cogn Neurosci. 2017a Jul;8(3):141–143. doi: 10.1080/17588928.2016.1273892

Slotnick SD. Cluster success: fMRI inferences for spatial extent have acceptable false-positive rates. Cogn Neurosci. 2017b Jul;8(3):150–155. doi: 10.1080/17588928.2017.1319350

Slotnick SD. No convincing evidence the hippocampus is associated with working memory. Cogn Neurosci. 2023 Jan-Oct;14(3):96–106. doi: 10.1080/17588928.2023.2223919

Slotnick SD. (2025a, Jul 18). Cluster threshold beta. Retrieved from osf.io/3wf7b

Slotnick SD. (2025b, Jul 22). Cluster threshold gamma. Retrieved from osf.io/69qcm

Slotnick SD, Moo LR, Segal JB, Hart J Jr. Distinct prefrontal cortex activity associated with item memory and source memory for visual shapes. Brain Res Cogn Brain Res. 2003 Jun;17(1):75–82. doi: 10.1016/s0926-6410(03)00082-x

Spets DS, Fritch HA, Thakral PP, Slotnick SD. High confidence spatial long-term memories produce greater cortical activity in males than females. Cogn Neurosci. 2021 Jul-Oct;12(3-4):112–119. doi: 10.1080/17588928.2020.1807924. Epub 2020 Aug 26. PMID: 32845219.

Spets DS, Karanian JM, Slotnick SD. False memories activate distinct brain regions in females and males. Neuroimage Rep. 2021 Aug 5;1(4):100043. doi: 10.1016/j.ynirp.2021.100043. PMID: 40568448; PMCID: PMC12172779.

Spets DS, Slotnick SD. Similar patterns of cortical activity in females and males during item memory. Brain Cogn. 2019 Oct;135:103581. doi: 10.1016/j.bandc.2019.103581. Epub 2019 Jul 10. PMID: 31301590.

Spets DS, Jeye BM, Slotnick SD. Different patterns of cortical activity in females and males during spatial long-term memory. Neuroimage. 2019 Oct 1;199:626–634. doi: 10.1016/j.neuroimage.2019.06.027. Epub 2019 Jun 14. PMID: 31207340.

Wagenmakers EJ, Love J, Marsman M, Jamil T, Ly A, Verhagen J, Selker R, Gronau QF, Dropmann D, Boutin B, Meerhoff F, Knight P, Raj A, van Kesteren EJ, van Doorn J, Šmíra M, Epskamp S, Etz A, Matzke D, de Jong T, van den Bergh D, Sarafoglou A, Steingroever H, Derks K, Rouder JN, Morey RD. Bayesian inference for psychology. Part II: Example applications with JASP. Psychon Bull Rev. 2018 Feb;25(1):58–76. doi: 10.3758/s13423-017-1323-7

